# A hyperthermoactive-Cas9 editing tool reveals the role of a unique arsenite methyltransferase in the arsenic resistance system of *Thermus thermophilus* HB27

**DOI:** 10.1101/2021.07.12.452139

**Authors:** Giovanni Gallo, Ioannis Mougiakos, Mauricio Bianco, Miriam Carbonaro, Andrea Carpentieri, Anna Illiano, Pietro Pucci, Simonetta Bartolucci, John van der Oost, Gabriella Fiorentino

**Affiliations:** Department of Biology, University of Naples Federico II, Via Cinthia 21, 80126 Napoli, Italy; Laboratory of Microbiology, Wageningen University & Research, Stippeneng 4, 6708WE Wageningen, The Netherlands; Department of Chemical Sciences, University of Naples Federico II, Via Cinthia 21, 80126 Napoli, Italy; Consiglio Nazionale delle Ricerche CNR, Institute of Polymers, Composites and Biomaterials (IPCB), via Campi Flegrei, 34, 80078 Pozzuoli, Na, Italy; Helmholtz Institute for RNA-based Infection Research (HIRI), Helmholtz-Centre for Infection Research (HZI), 97080 Würzburg, Germany; CEINGE-Biotecnologie avanzate, Via Comunale Margherita, 484-538, 80131 Napoli NA; Task Force on Microbiome Studies, University of Naples Federico II, Naples, Italy

**Keywords:** Arsenic resistance, thermoresistance, extremophiles, *Thermus thermophilus*, CRISPR-Cas9 genome editing, genetic tool, bioreporter

## Abstract

Arsenic detoxification systems can be found in a wide range of organisms, from bacteria to man. In a previous study, we discovered an arsenic-responsive transcriptional regulator in the thermophilic bacterium *Thermus thermophilus* HB27 (*Tt*SmtB). Here, we characterize the arsenic resistance system of *T. thermophilus* in more detail. We employed *Tt*SmtB-based pull-down assays with protein extracts from cultures treated with arsenate and arsenite to obtain an S-adenosyl-Lmethionine (SAM)-dependent arsenite methyltransferase (*Tt*ArsM). *In vivo* and *in vitro* analyses were performed to shed light on this new component of the arsenic resistance network and its peculiar catalytic mechanism. Heterologous expression of *TtarsM* in *Escherichia coli* resulted in arsenite detoxification at mesophilic temperatures. Although *Tt*ArsM does not contain a canonical arsenite binding site, the purified protein does catalyze SAM-dependent arsenite methylation with formation of monomethylarsenite (MMAs) and dimethylarsenite (DMAs). In addition, *in vitro* analyses confirmed the unique interaction between *Tt*ArsM and *Tt*SmtB. Next, a highly efficient ThermoCas9- based genome-editing tool was developed to delete the *Tt*ArsM-encoding gene on the *T. thermophilus* genome and to confirm its involvement in the arsenite detoxification system. Finally, the *TtarsX* efflux pump gene in the *T. thermophilus* Δ*TtarsM* genome was substituted by a gene encoding a stabilized yellow fluorescent protein (sYFP) to create a sensitive genome-based bioreporter system for the detection of arsenic ions.

**Importance:** We here describe the discovery of an unknown protein by using a proteomic approach with a functionally related protein as bait. Remarkably, we successfully obtained a novel type of enzyme through the interaction with a transcription regulator, controlling the expression of this enzyme. Employing this strategy, we isolated *Tt*ArsM, the first thermophilic prokaryotic arsenite methyltransferase, as a new enzyme of the arsenic resistance mechanism in *T. thermophilus* HB27. The atypical arsenite binding site of *Tt*ArsM categorizes the enzyme as the first member of a new arsenite methyltransferase type, exclusively present in the *Thermus* genus. The enzyme methylates arsenite producing MMAs and DMAs. Furthermore, we developed an hyperthermophilic Cas9-based genome-editing tool, active up to 65°C. The tool allowed us to perform highly efficient, marker-free modifications (either gene deletion or insertion) in the *T. thermophilus* genome. With these modifications, we confirmed the critical role of *Tt*ArsM in the arsenite detoxification system and developed a sensitive *whole cell* bioreporter for arsenic ions. We anticipate that the developed tool can be easily adapted for editing the genomes of other thermophilic bacteria, significantly boosting fundamental and metabolic engineering in hyperthermophilic microorganisms.

## Introduction

Arsenic is the most abundant environmental toxic element which enters the biosphere mainly from geochemical and (to a lesser extent) anthropogenic sources such as herbicides, growth promoters for livestock, and industrial activities (1). Arsenic has two relevant oxidation states, trivalent arsenite As(III) and pentavalent arsenate As(V). Methylated arsenicals include mono- (MAs), di- (DMAs), and tri- (TMAs) methylated forms. In general, trivalent states are more toxic than the pentavalent ones, and TMAs is more toxic than inorganic arsenite. Although arsenic is not beneficial for life, it can enter cells through transporters such as aquaglyceroporins. Hence, arsenic detoxification systems can be found in a wide range of organisms, from bacteria to humans. Arsenic resistance genes (*ars*) include genes encoding efflux transporters, redox enzymes, methyltransferases, transcriptional repressors and biosynthetic pathways for arsenosugars and arsenolipids (2, 3). The identification and characterization of these pathways have attracted the attention of fundamental, evolutionary, and biotechnological research (4–6).

Microorganisms have been exposed to arsenic since the origin of life and consequently have evolved arsenic resistance systems, encoded by genes generally clustered in operons (7). The organization and number of the operons of arsenic resistance genes are highly variable between different species (8, 9), reflecting differences in the level of arsenic resistance. The key players of arsenic detoxification are: (i) arsenate reductases (ArsC) that reduce intracellular arsenate to arsenite (10), (ii) efflux permeases responsible for arsenite transport outside the cell (11), and (iii) transcriptional repressors that are generally metalloregulatory proteins of the ArsR/SmtB family (12). In addition, arsenite can be methylated by arsenite *S*-adenosylmethionine methyltransferases (ArsM) into MMAs, DMAs and TMAs (13), after which they can passively leave the cell or be extruded by methylarsenite-specific efflux permease (ArsP) (14).

Based on recent molecular clock analyses, it has been concluded that arsenite efflux and arsenite methylation represented the core of microbial arsenic resistance systems before the rise of atmospheric oxygen (15). In such primordial anoxic environments, methyl-arsenicals could also have the function as antibiotics against competitor microbes; after the rise of atmospheric oxygen, the ArsM enzymes did become primary components of the arsenic detoxification machinery; nevertheless, in some microorganisms, they maintained their antibiotic activity (16, 17).

Hydrothermal hot springs, which can be considered environments with conditions similar to niches of primordial Earth, may contain high amounts of arsenic. In these cases, these hot springs are niches for arsenic-tolerant microorganisms, which play a critical role in the global arsenic biogeochemical cycle (18). Although the resistance mechanisms to inorganic arsenic have been studied in many microorganisms (19), the contribution of organo-arsenical biotransformation in extreme environments is still at a stage of infancy. In this regard, only two algal thermoactive ArsM enzymes have been characterized to date (20).

The thermophilic bacterium *Thermus thermophilus* HB27, originally isolated from a volcanic hot spring in Japan (21), has an unusual genetic organization of its machinery to cope with arsenic toxicity. The currently identified arsenic resistance genes are randomly scattered in its genome (22), complicating the identification of all the genes involved. In previous studies, we elucidated some components of the arsenic resistance system of *T. thermophilus* HB27. We identified and characterized *Tt*SmtB, the metalloregulatory transcriptional repressor that is responsible for the regulation of the arsenic detoxification system. *Tt*SmtB recognizes and firmly binds to operator sequences in the promoter regions of the arsenite efflux gene (*TtarsX*) (23) and the arsenate reductase gene (*TtarsC*) (24, 25), efficiently repressing their transcription in the absence of arsenic ions. *Tt*SmtB and *Tt*ArsX are also involved in cadmium sensing and export, respectively (23). In two *T. thermophilus* HB27 deletion mutant strains (Δ*TtarsX* and Δ*TtsmtB*), tolerance to arsenate, arsenite and cadmium was significantly reduced compared to the wild type strain. Although these analyses confirmed the involvement of *Tt*ArsX and *Tt*SmtB in the promiscuous resistance mechanism, the mutant strains could still grow at concentrations of arsenic up to 3 mM (22, 23). Notably, the genome of *T. thermophilus* HB27 is not predicted to express arsenite methyltransferases or arsenite oxidases, suggesting the existence of unidentified component(s) of the *T. thermophilus* HB27 arsenic resistance system that cannot be predicted by *in silico* approaches, highlighting the need to employ an alternative experimental identification method.

Since members of the ArsR/SmtB family are a group of homodimeric proteins with a common HTH-winged helix DNA binding domain and heterogeneous metal-binding domain architectures and interaction modes (26), we hypothesized that *Tt*SmtB could even form protein interactions with unknown, functionally-related protein partners.

In this study, using an integrated proteomic, biochemical and genetic approach, we provide a gain of insight into the arsenic resistance system of *T. thermophilus* HB27. We report the discovery of the first *T. thermophilus* HB27 arsenite methyltransferase, *Tt*ArsM. Moreover, we describe the development of a highly efficient, markerless Cas9-based genome-editing tool at temperatures up to 65°C. Using this ThermoCas9 system, we demonstrated the *in vivo* involvement of *Tt*ArsM in arsenite detoxification. The newly developed genome editing tool was further validated constructing a very sensitive *whole cell* bioreporter system in which the *TtarsX* efflux transporter gene was substituted by a gene encoding a thermo-adapted superfolder yellow fluorescent protein (*syfp*) (27).

## Results

### Exploring the protein-protein interactions of *Tt*SmtB

A combined comparative and functional proteomic approach was employed to identify putative *Tt*SmtB interacting proteins with a role in arsenite metabolism/detoxification. Purified recombinant His-tagged *Tt*SmtB was bound to a Ni^2+^/NTA resin for protein pull-down assays using *T. thermophilus* HB27 cell-free extracts (CFE) from cultures exposed either to arsenite or arsenate, or untreated CFE cultures used as control. SDS-PAGE separation of the pulled-down proteins eluted with 0.5 M imidazole, followed by LC-electrospray ionization (ESI)-MS/MS (28) and comparative analysis of the acquired data resulted in the identification of 51 cytosolic proteins that interact with *Tt*SmtB (Table S1). Only five of these proteins are simultaneously present in CFE from cultures exposed to arsenite and arsenate but not in control CFE from the non-exposed cultures. Amongst these proteins, TTC0109 (Accession No. AAS80457, UniProt code Q72LF0) was predicted to be involved in the arsenic detoxification system (based on homology to annotated ArsR family transcriptional regulators) and to contain a C-terminal SAM-dependent methyltransferase domain (based on homology to annotated methyltransferase domain-containing proteins), suggesting a role in arsenic methylation. To date, there are no annotated arsenite SAM-dependent methyltransferases in the genome of *T. thermophilus* HB27 or the genomes of other thermophilic bacteria. Hence, we selected TTC0109 for further investigation as a potential novel arsenite methyltransferase.

### Bioinformatic analysis of TTC0109

BlastP analysis of TTC0109 translated sequence with sequenced microbial genomes and evolutionary analysis conducted with MEGA X, demonstrated that TTC0109 is highly conserved among the members of the *Thermus* genus (Fig. 1 A). Moreover, multiple sequence alignment of TTC0109 with characterized prokaryotic arsenite methyltransferases (Fig. 1 B) showed that all the aligned proteins contain a typical Rossman fold (29). This fold contains a GxGxG motif in a loop region, which presumably interacts with the carboxypropyl moiety of SAM, and a highly conserved aspartic acid residue at the end of the β2 strand which forms hydrogen bonds with the ribose hydroxyls of the cofactor (30). In the case of TTC0109, the predicted GxGxG motif is composed of G114, T115, G116, T117, G118 residues and the conserved aspartic residue is D135 (29, 30). On the other hand, the alignment shows that TTC0109 greatly differ from characterized ArsM proteins in the remaining sequence (Fig. 1 B); nonetheless, if TTC0109 is an arsenite methyltransferase, it would be evolutionarily distant from other archaeal and bacterial arsenite methyltransferases, as shown in the phylogenetic tree (Fig. 1 A), and therefore it could belong to a new type of arsenite methyltransferase. Additionally, all the known arsenite methyltransferases, including those in the alignment, possess at least two, usually three, cysteines that are responsible for the binding of arsenite and its subsequent methylation (16, 31). TTC0109 contains a single cysteine residue at position 77, which is perfectly conserved in all the sequences analyzed (Fig. 1 B); hence, TTC0109 could be an arsenite methyltransferase with a distinct reaction mechanism.

**Fig. 1.**
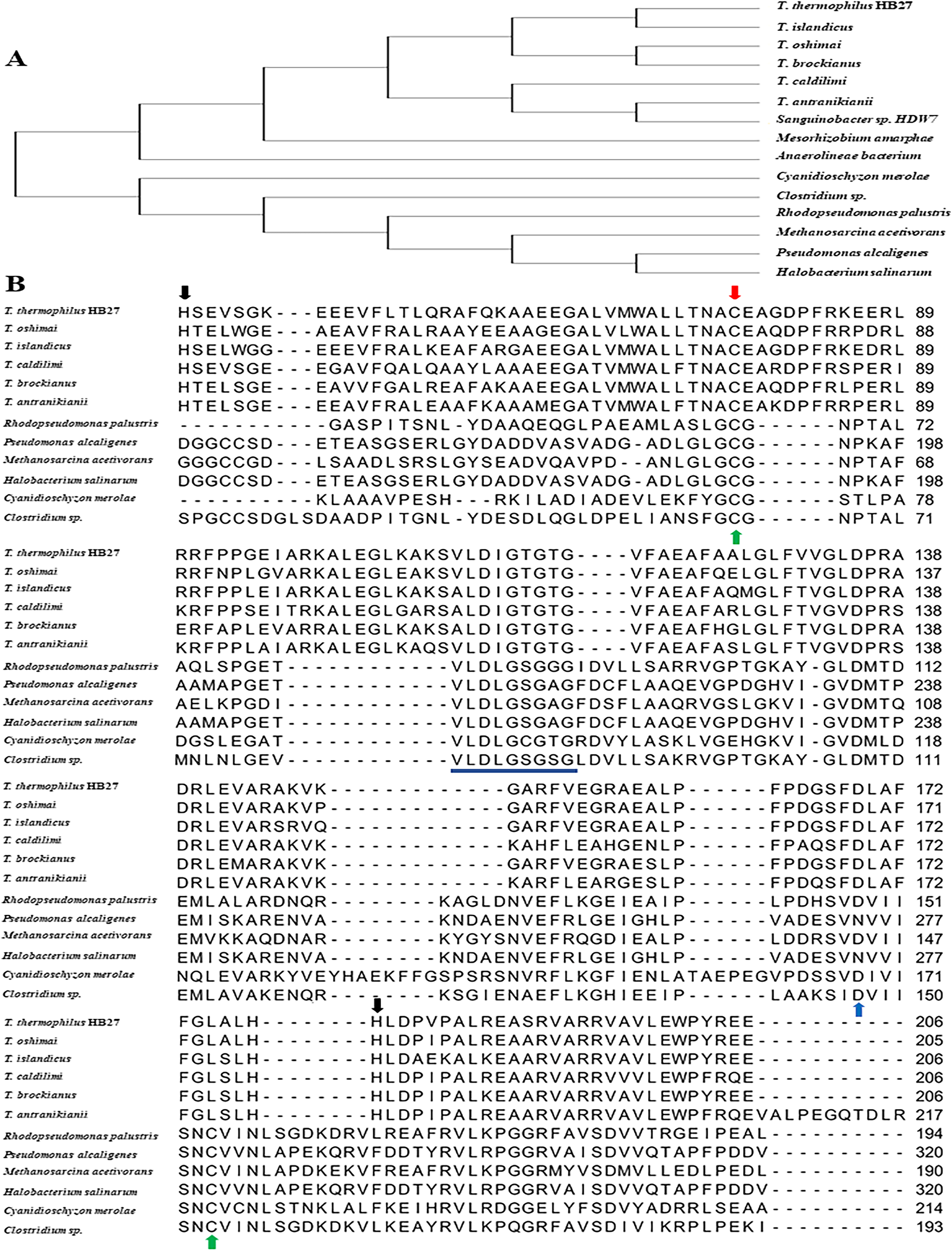
**A) Phylogenetic tree of archaeal and bacterial arsenite methyltransferases, SAM-dependent methyltransferases and methyltransferase domain containing proteins.** The aminoacidic sequences used for the construction of the phylogenetic tree are: *Tt*ArsM from *T. thermophilus* HB27; SAM-dependent methyltransferase from five members of the *Thermus* genus (*islandicus, caldilimi, antranikianii, oshimai* and *brockianus*), *Anaerolineae bacterium* and *Mesorhizobium amorphae*; arsenite methyltransferase from *Clostridium* sp BMX (53)*, Methanosarcina acetivorans, Rhodopseudomonas palustris* (13)*, Pseudomonas alcaligenes* (36) *Halobacterium salinarum* (54) *Cyanidioschyzon merolae* (45) and a methyltransferase domain-containing protein from *Sanguinobacter* sp. **B) Multiple sequence alignment of hypothetical and functionally characterized arsenite methyltransferases (ArsM) with *T. thermophilus* HB27 *Tt*ArsM.** The partial alignment includes 5 members of the *Thermus* genus (*islandicus, caldilimi, antranikianii, oshimai* and *brockianus*) (98% identity to *Tt*ArsM, sequence aligned from amino acid 40 to 89), *Clostridium* sp BMX (53) (28% identity to *Tt*ArsM, from 22 to 193)*, R. palustris* (13) (32% identity to *Tt*ArsM, from 23 to 168), *M. acetivorans* (37) (29% identity to *Tt*ArsM, from 19 to 190), *P. alcaligenes* (36) (31% identity to *Tt*ArsM, from 149 to 320), *Cyanidioschyzon merolae* (45) (27.7% identity to *Tt*ArsM, from 29 to 214) and *H. salinarum* (54) (25% identity to *Tt*ArsM, from 58 to 294). Red arrow indicates the catalytic cysteine, conserved in characterized ArsM and *Tt*ArsM; green arrows indicate two catalytic cysteines conserved in characterized ArsM, but not in *Tt*ArsM; blue arrow indicates the conserved aspartic acid; the SAM binding domain, which is part of the typical Rossman fold, is underlined in blue. The two histidines of *Tt*ArsM predicted to interact with arsenite are indicated by black arrows.

Reasoning that the TTC0109 structure could provide more information regarding the function of TTC0109, we generated a structural model of the protein and performed molecular docking with arsenite and SAM (Fig. S1). The obtained model predicts that TTC0109 forms homodimers via its N-terminal moiety; molecular docking highlighted that arsenite could be coordinated by two histidines, H40 and H179, while C77 interacts with the methyl group of SAM (Fig. S1 A).

Although H40 and H179 residues are not conserved in characterized ArsM proteins, they are maintained at an identical position in the translated genomes of all *Thermus* species (five of them are shown in Fig. 1B); moreover, H40 is encompassed in a sequence motif (34-<u>YR</u>VFPT**<u>H</u>**S<u>E</u>-42) that shares 45% identity (underlined) with a sequence motif (101-<u>YR</u>LADR**<u>H</u>**V<u>E</u>-109) at the C-terminus of the *Tt*SmtB metal-binding site (32), strengthening the hypothesis that H40 could be involved in arsenite binding, and suggesting an evolutionary connection between the two proteins.

We proceeded with the generation of structural models of mutant proteins in which the amino acids H40 or H179 were replaced with alanine residues producing TTC0109 H40A, TTC0109 H179A and C77 replaced with a serine residue producing TTC0109 C77S (Fig. S1 B, C, D). The predicted models showed that the substitution of either H40 or H179 with an alanine residue altered the 3D structure of TTC0109, whereas the effect of the cysteine to serine substitution had only a minimal effect (Fig. S1 C, D), supporting the hypothesis that this residue could have a functional role.

### TTC0109 is a novel arsenite methyltransferase

A recombinant His-tagged version of TTC0109 was produced and purified to homogeneity from *E. coli* BL21-CodonPlus (DE3)-RIL cells transformed with pET30b(+)*/TtarsM* (predicted mass: 29.1 kDa) (Fig. S2 A). Gel filtration chromatography analysis agreed with the *in silico* predicted dimeric configuration of the protein, showing that the homodimer has a mass of approximately 64.5 kDa (Fig. S2 B). To determine whether TTC0109 had arsenite methyltransferase activity, a coupled spectrophotometric enzymatic assay based on the formation of S-adenosylhomocysteine (SAH) from SAM after transfer of methyl-group(s) on the substrate was employed (33, 34). In this case, the acceptor of methyl groups was As(III). SAH is degraded by SAH nucleosidase into S-ribosylhomocysteine and adenine; adenine deaminase acts on adenine producing hypoxanthine, which is converted into urate and hydrogen peroxide (H_2_O_2_) by xanthine oxidase. The rate of production of H_2_O_2_ is measured by an increase in absorbance at 510_nm_ with the help of the colorimetric reagent 3,5-dichloro-2-hydroxybenzensulfonic acid (DHBS). Then, arsenite methyltransferase activity was assayed following the increase in absorbance at 510_nm_ (35). Preliminary assays were set up to assess the thermal stability of the different components, and consequently the optimal assay temperature; afterwards, the saturating concentrations of SAM and arsenite were determined. Therefore, the optimal assay conditions resulted: 50°C, 200 μM arsenite, 800 μM SAM and 3.1 μM of TTC0109. Under these conditions, the specific arsenite methyltransferase activity of TTC0109 was 4.5 mU/mg (Fig. 2 A). For this reason, hereafter, the TTC0109 protein will be denoted as *Tt*ArsM.

**Fig. 2.**
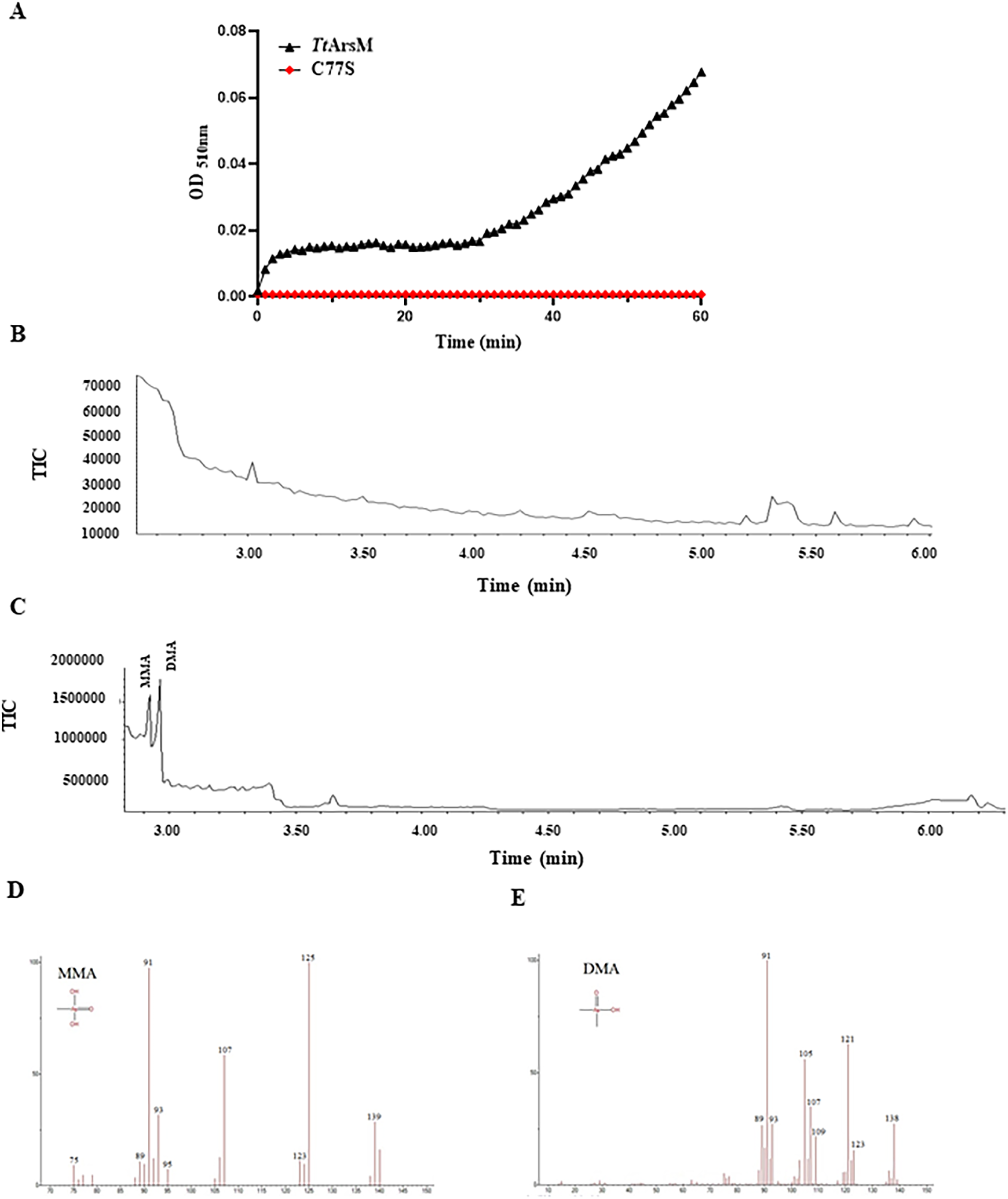
***In vitro* assessment of methyltransferase activity. A) Arsenite methylation assays.** The enzyme coupled colorimetric assay was carried out in continuous in the presence of 3.1 μM of recombinant *Tt*ArsM (black curve) or *Tt*ArsM C77S (red curve), 800 µM SAM and 200 µM As(III) at 50°C. The absorbance of the reaction mixture was recorded every minute for a total of 1 hour. The graph represents the average of three independent experiments, each performed in triplicate. **B) Products of As(III) methylation by purified *Tt*ArsM.** Arsenic species were analyzed by GC-MS. Each assay contained 10 µM *Tt*ArsM, 250 µM As(III), 6 mM GSH and 1 mM SAM, incubated at 65°C for 24 h. In **B)** and **C)**, the chromatograms recorded between 2 and 6 min for negative control and the enzyme mixture. **D** and **E** report the fragmentation spectra for MMAs and DMAs.

In order to characterize which products are formed upon As(III) methylation by *Tt*ArsM, we incubated 10 µM of *Tt*ArsM with As(III), GSH and SAM, at 65°C for 24 h; the mixture was then solubilized and analyzed by GC-MS. The results obtained are shown in Figure 2 C. Two sharp peaks at 2.92 min and 2.96 min attributed to MMAs and DMAs, respectively, are only visible in the enzyme mixture, as no peaks were recorded for the control sample (Fig. 2 B). The fragmentation spectra of MMAs and DMAs are reported in Fig 2 D and E. Peak areas corresponding to MMAs and DMAs were manually integrated, and the results are summarized in Figure S2 C.

The results of *in vitro* assays using purified *Tt*ArsM protein confirmed the ability of the protein to methylate As(III) producing mono and dimethylated arsenic, the latter being the primary product. The oxidation state of the products could not be determined because the reactions were terminated with H_2_O_2_, which oxidized all arsenicals to pentavalent states.

Since the *in silico* predictions of *Tt*ArsM led to the hypothesis that C77, H40 and H179 were catalytic amino acids, three mutated versions of *TtarsM,* namely *TtarsM* C77S, *TtarsM* H40A and *TtarsM* H179A, were constructed, expressed in *E. coli* BL21-CodonPlus (DE3)-RIL cells and the corresponding *Tt*ArsM mutants were purified (Fig S3). Although the expression levels of the three mutant proteins are comparable (Fig S3), it was not possible to perform *in vitro* characterization of purified *Tt*ArsM H40A and *Tt*ArsM H179A, which precipitated in solution after purification; this phenomenon is probably due to protein instability, thus indicating the importance of these amino acids for *Tt*ArsM structure. On the other hand, the purification of soluble *Tt*ArsM C77S protein was possible, albeit with a lower yield compared to the wild type *Tt*ArsM. Nonetheless, using the previously mentioned coupled assay, this mutant enzyme did not show any *in vitro* arsenite methyltransferase activity, confirming that C77 residue plays a role in *Tt*ArsM activity (Fig. 2 A). These *in vitro* results demonstrated that *Tt*ArsM has an arsenite methyltransferase activity and its distinct active site suggests a novel reaction mechanism when compared to other characterized arsenite methyltransferases (13, 36, 37).

### *Tt*SmtB interacts with *Tt*ArsM and binds to its promoter

This is the first study to report the protein-protein interaction of an ArsR/SmtB transcriptional regulator with a member of the arsenic detoxification system as identified by pull-down and mass spectrometry. For this reason, we decided to confirm the physical interaction between *Tt*SmtB and *Tt*ArsM and to investigate the effect of different metals on *Tt*SmtB:*Tt*ArsM interaction. A Co-IP assay was carried out upon incubation of purified *Tt*ArsM and *Tt*SmtB either in the presence or in the absence of arsenite, arsenate, cadmium and antimony. The first three ions are *Tt*SmtB effectors as their interaction weakens the binding to target promoters, while antimony has no effect on DNA recognition (32). Immunoprecipitation with anti-*Tt*SmtB antibodies, followed by detection of the His-tagged *Tt*ArsM by anti-His-tag antibodies, showed that the two proteins interact and form a complex in the absence of arsenite and arsenate, confirming the existence of physical interaction between them (Fig. 3). No band was detected when the immunoprecipitation was carried out with the unrelated control protein *Tt*GalA (38). Increasing arsenate and arsenite concentrations negatively affected the stability of *Tt*SmtB:*Tt*ArsM complex; in fact, densitometric analysis of the Western-blot revealed up to a 3-fold decrease in the intensity of the band corresponding to the complex (at 1:100 protein:arsenic ratio) in the presence of both arsenate (Fig. 3 A) and arsenite (Fig. 3 B). Interestingly, the presence of cadmium had the opposite effect, enhancing by up to 2-fold the band intensity (Fig. 3 C), suggesting that the interaction of this metal with the complex occurs with a different mechanism. Finally, the presence of antimony had a negligible effect on complex stability, in agreement with previous data showing that this metal ion is not an effector for *Tt*SmtB (32) (Fig. 3 D).

**Fig. 3.**
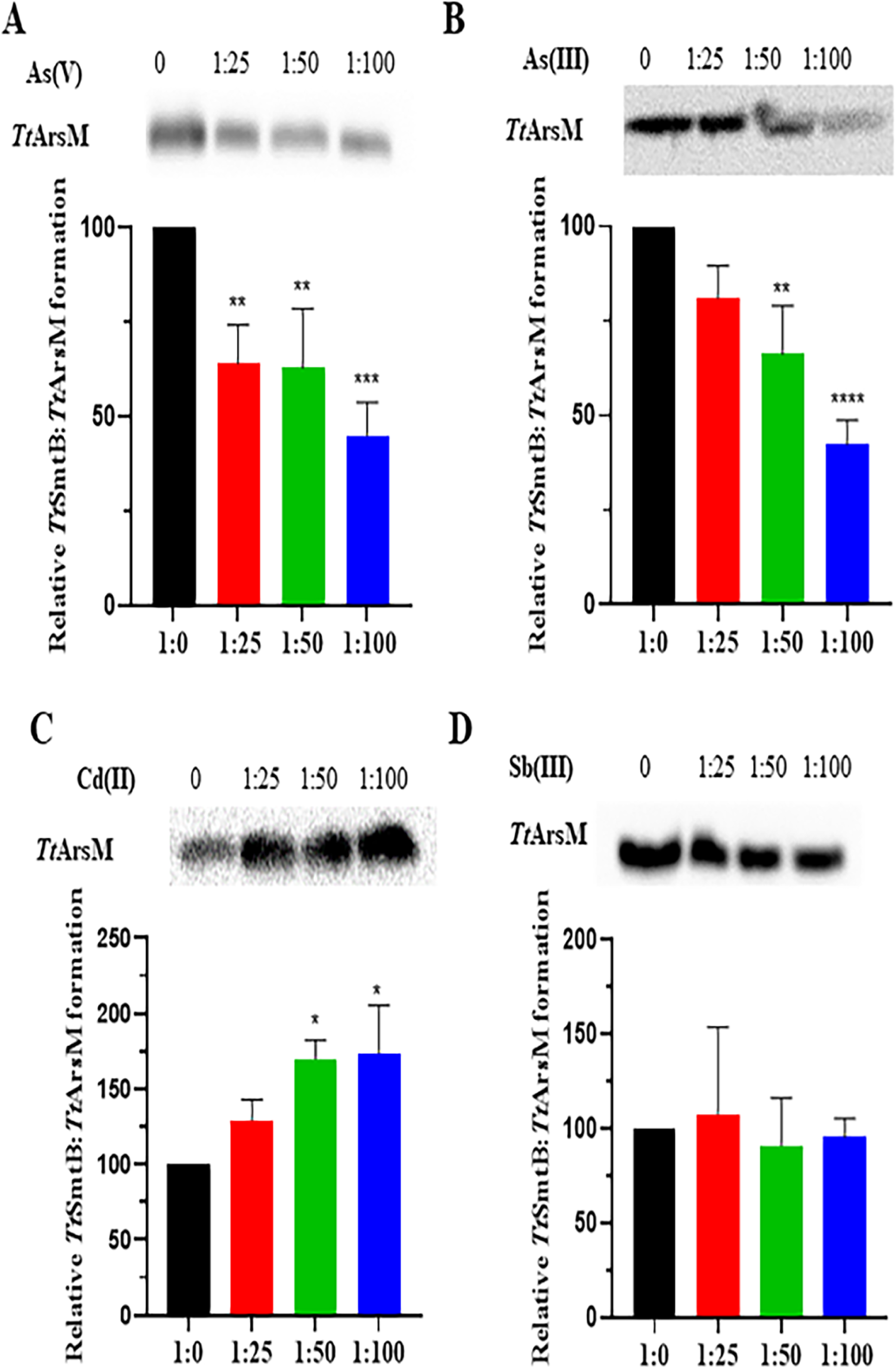
***Tt*SmtB:*Tt*ArsM interaction in the presence of heavy metal ions.** CoIp of *Tt*SmtB:*Tt*ArsM complex with increasing concentrations of **(A)** arsenate, **(B)** arsenite, **(C)** cadmium and **(D)** antimony. Complexes were immunoprecipitated with anti-*Tt*SmtB antibodies and revealed through Western-blot using anti-His antibodies against the His-tag of *Tt*ArsM. Below: densitometric analysis of blots of *Tt*SmtB:*Tt*ArsM complex. The intensity of the unchallenged complex was used as a reference. Average values from three biological replicates are shown, with error bars representing standard deviations. Statistical analysis was performed using one-way ANOVA; significant differences are indicated as: * p < 0.05, ** p < 0.01, *** p < 0.001, **** p < 0.0001.

Since *Tt*SmtB is the transcriptional repressor of the genes involved in arsenic and cadmium resistance in *T. thermophilus* HB27 (22, 23), we hypothesized that it could also regulate *TtarsM* transcription. Sequence analysis of *TtarsM* promoter (*p_arsM_*), a 108 bp-long region upstream of *TtarsM* and encompassing the translation start codon revealed the presence of an inverted repeat region [GAAC(N14)CTTG] between positions -6 and -27 upstream of the start codon. The sequence overlaps -10 and -35 putative basal promoter region, is 100% identical to the *TtarsX* operator recognized by *Tt*SmtB and matches the consensus binding sites of ArsR/SmtB proteins (39). Hence, we performed EMSA to investigate the capacity of purified *Tt*SmtB to bind to the promoter region of *TtarsM*. *Tt*SmtB binds to *p_arsM_* in a concentration-dependent manner, as shown by the gradual formation of lower mobility complexes and the gradual decrease of residual unbound DNA (Fig. 4 A, lanes 2-5); at 10 μM protein, the complex hardly enters the gel, suggesting the formation of multiple dimers associated with target DNA (Fig. 4 A lane 6). This observation suggests that by interacting with the regulatory region *Tt*SmtB controls *TtarsM* transcription in a way comparable to that already reported for other arsenic resistance genes, i.e. the arsenate reductase, the arsenite/cadmium efflux transporter and itself (22, 23).

**Fig. 4.**
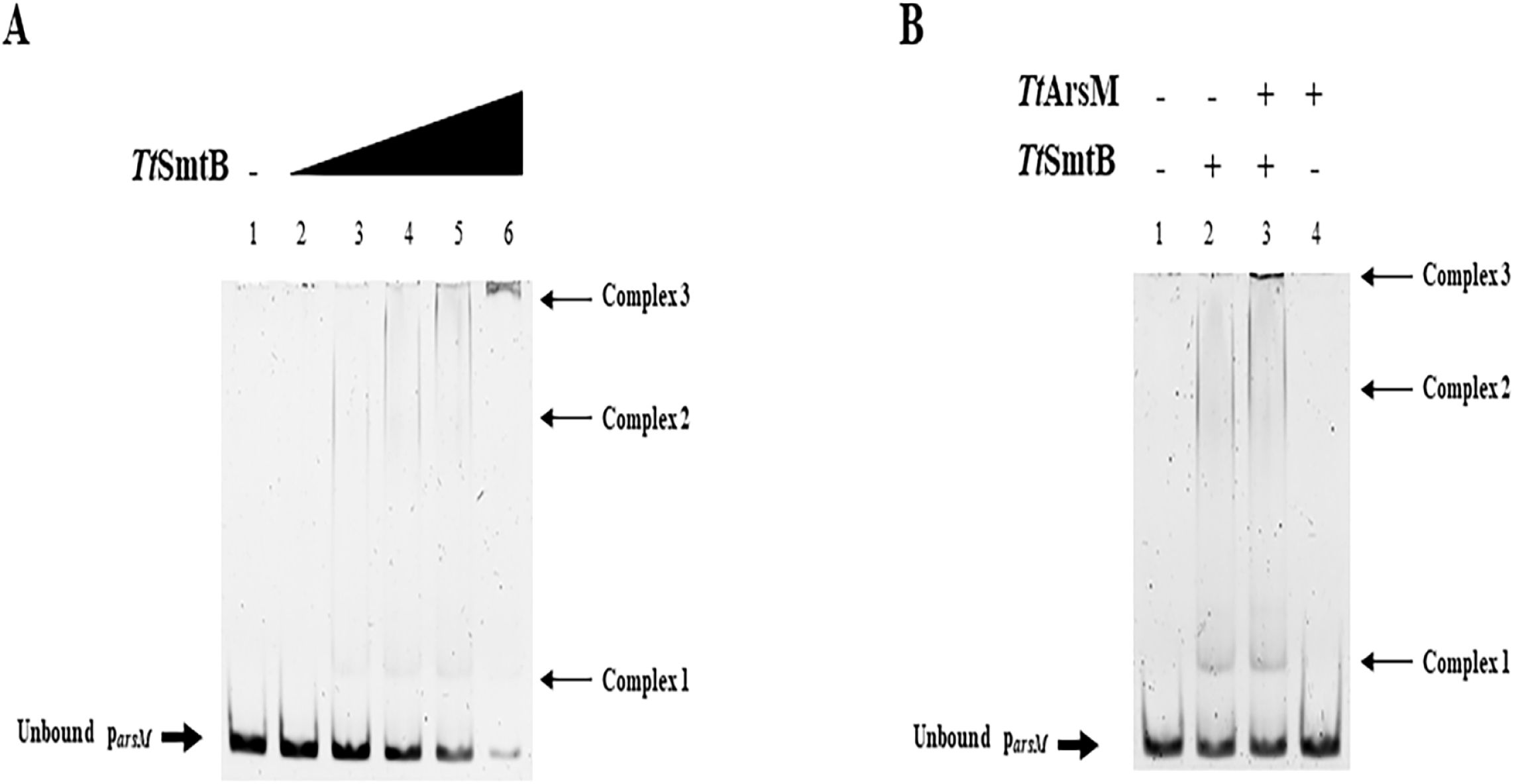
***Tt*SmtB:*p_arsM_* interaction. (A)** Interaction of *p_arsM_* in the presence of increasing concentration of *Tt*SmtB. Lane 1: negative control; Lane 2: 1 µM *Tt*SmtB; Lane 3: 2 µM *Tt*SmtB; Lane 4: 3 µM *Tt*SmtB; Lane 5: 5 µM *Tt*SmtB; Lane 6: 10 µM *Tt*SmtB. **(B)** EMSA with *p_arsM_*, in presence of 3 µM *Tt*SmtB and 3 µM *Tt*ArsM.

Since the existence of a physical interaction between *Tt*SmtB and *Tt*ArsM was established, we asked whether *Tt*ArsM influenced *Tt*SmtB interaction with *p_arsM._* Therefore, we pre-incubated 3 μM of both proteins before performing an EMSA in the same conditions described above. Interestingly, when the two proteins are co-incubated, shifted bands can be observed (Fig 4 B lane 3, complex 3) corresponding to complexes of higher molecular weight in comparison to those generated or not by *Tt*SmtB or *Tt*ArsM, respectively (Fig 4 B lane 2 complex 2 and lane 4); this analysis indicates that *Tt*SmtB:*Tt*ArsM multimeric complexes bind to the promoter and suggests that *Tt*SmtB:*Tt*ArsM protein-protein interaction may function in either transcriptional and post-transcriptional control. Notably, very few studies in bacteria report protein-protein interactions of transcriptional regulators with the product of the genes they regulate (40, 41).

### *In vivo* activity of *Tt*ArsM and its catalytic mutants in *E. coli*

Aiming to explore the role of *Tt*ArsM in arsenite resistance *in vivo*, we challenged *E. coli* BL21-CodonPlus (DE3)-RIL strains transformed with plasmids expressing *Tt*ArsM and its catalytic mutants (*Tt*ArsM C77S, *Tt*ArsM H40A and *Tt*ArsM H179A) to grow in the presence of arsenite. Each recombinant strain was grown in the presence of different arsenite concentrations for 24-hours to determine the minimal inhibitory concentration (MIC) towards the metal ion.

*Tt*ArsM-expressing strain appeared to be more resistant to arsenite than the control strain (MIC 6 mM and 4.5 mM, respectively). Additionally, the strains expressing mutated *Tt*ArsM were inhibited by the presence of arsenite to the same extent as the control strain (Fig. 5). This shows that the heterologous expression of *Tt*ArsM in *E. coli* increases arsenite resistance even at mesophilic temperatures, indicating the role of *Tt*ArsM in arsenite detoxification. Moreover, the result obtained with the mutant strains demonstrates the role of C77, H40 and H179 in the catalytic function of *Tt*ArsM.

**Fig. 5.**
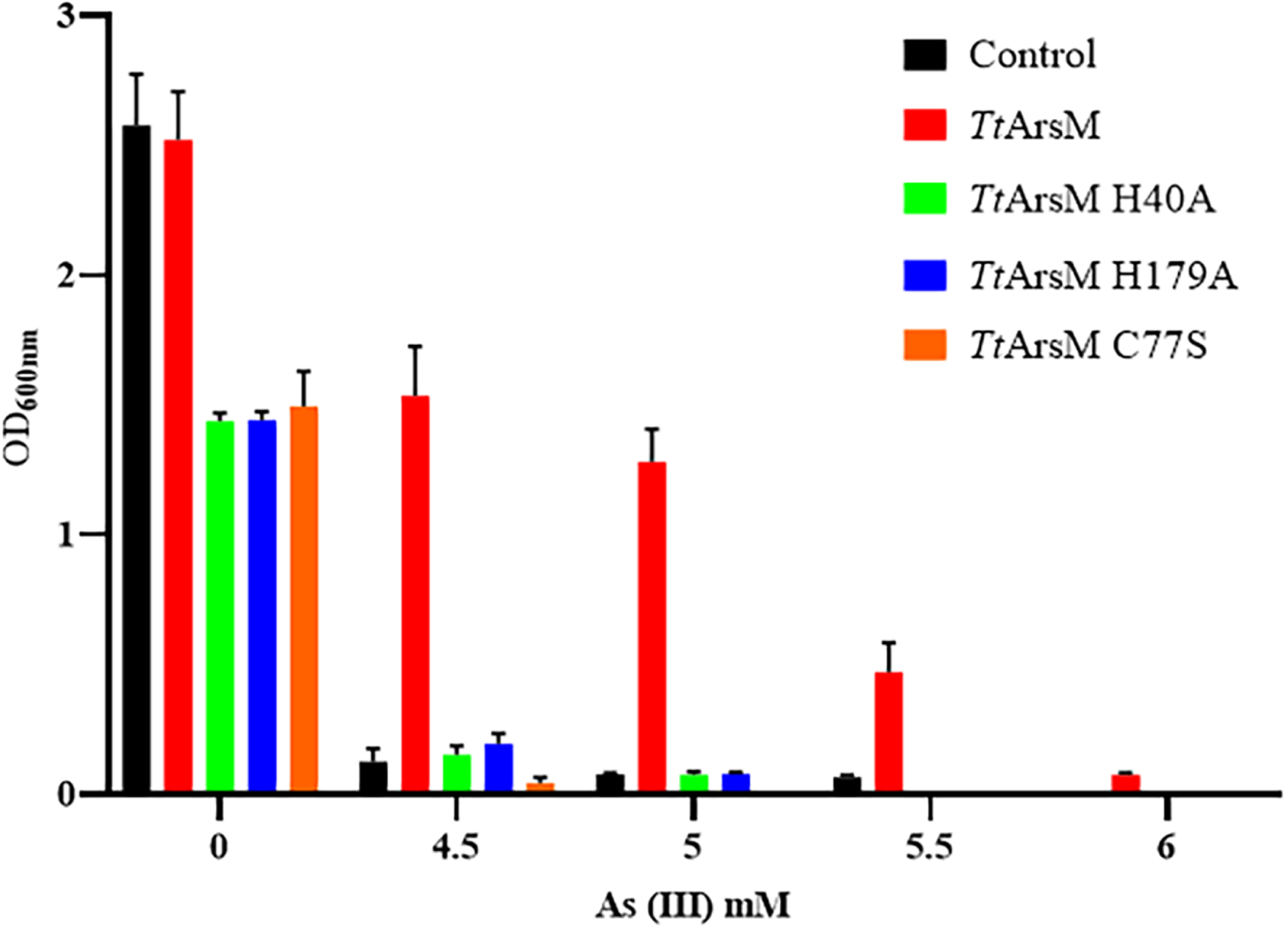
**Growth of *E. coli* BL21 strains expressing *Tt*ArsM and its mutants, in the presence of different arsenite concentrations, measured 24 hours post-inoculation.** The strains are: *E. coli* BL21/pET30b (black), *E. coli* BL21/pET30/*Ttars*M (red), *E. coli* BL21/pET30/ *Ttars*M H40A (green), *E. coli* BL21/pET30/*Ttars*M H179A (blue) and *E. coli* BL21/pET30/*TtarsM* C77S (orange). Average values from three biological replicates are shown, with error bars representing standard deviations.

### Developing a hyperthermoactive Cas9 editing tool

We further aimed to investigate *in vivo* the contribution of *Tt*ArsM to the arsenite detoxification mechanism via the deletion of the *TtarsM* gene from the *T. thermophilus* HB27 genome. Nonetheless, the currently available genome editing tool for *T. thermophilus* is time-consuming, not marker-free and not always efficient (42). For this purpose, we reasoned to develop a marker-free, plasmid-based, homologous recombination (HR) Cas9 counter-selection (CS) genome editing tool for *T. thermophilus* employing ThermoCas9, a thermotolerant and thermoactive Cas9 orthologue (43).

We initially evaluated the targeting efficiency of ThermoCas9 in *T. thermophilus* HB27. Therefore, a set of 3 vectors was constructed, namely pMK-ThermoCas9-NT, pMK-ThermoCas9-sp1 and pMK-ThermoCas9-sp2 (Fig. 6 A), by cloning into the pMK18 vector (44) (i) the codon-harmonized version of the *thermocas9* gene under the transcriptional control of the constitutive *nqo* promoter (42), and (ii) the sgRNA expressing module under the transcriptional control of the constitutive 16S rRNA promoter, either with a non-targeting/control spacer (NT: 5’- CTAGATCCGCAGTAACCCCATGG-3’) or with spacers that target the *TtarsM* gene (sp1: 5’-GGGCGTTGGTGATGTGGGCCCTC-3’ and sp2: 5’- CCACCTCCTCCTCCCGGTAAGGC-3’). The 3 vectors were used to transform *T. thermophilus* HB27, along with pMK-Pnqo-syfp vector (27) as transformation control. The cells were allowed to recover at 70°C, before being plated on selective agar plates and incubated overnight at 60°C, due to the sensitivity of pMK18 at temperatures above 65°C. The transformation efficiencies of pMK- ThermoCas9-sp1 and pMK-ThermoCas9-sp2 targeting vectors were significantly reduced compared to the transformation efficiency with the pMK- ThermoCas9-NT non-targeting vector (Fig. 6 B). Moreover, the transformation efficiency with the pMK-Pnqo-syfp vector was only slightly higher compared to the transformation efficiency with the pMK-ThermoCas9-NT vector (Fig. 6 B), which could be attributed to the significant size difference between the two vectors (7552 bp and 13554 bp, respectively). This result indicates that ThermoCas9 is expressed in *T. thermophilus* HB27 cells in an active and not toxic form, motivating the development of a ThermoCas9-based genome-editing tool.

**Fig. 6.**
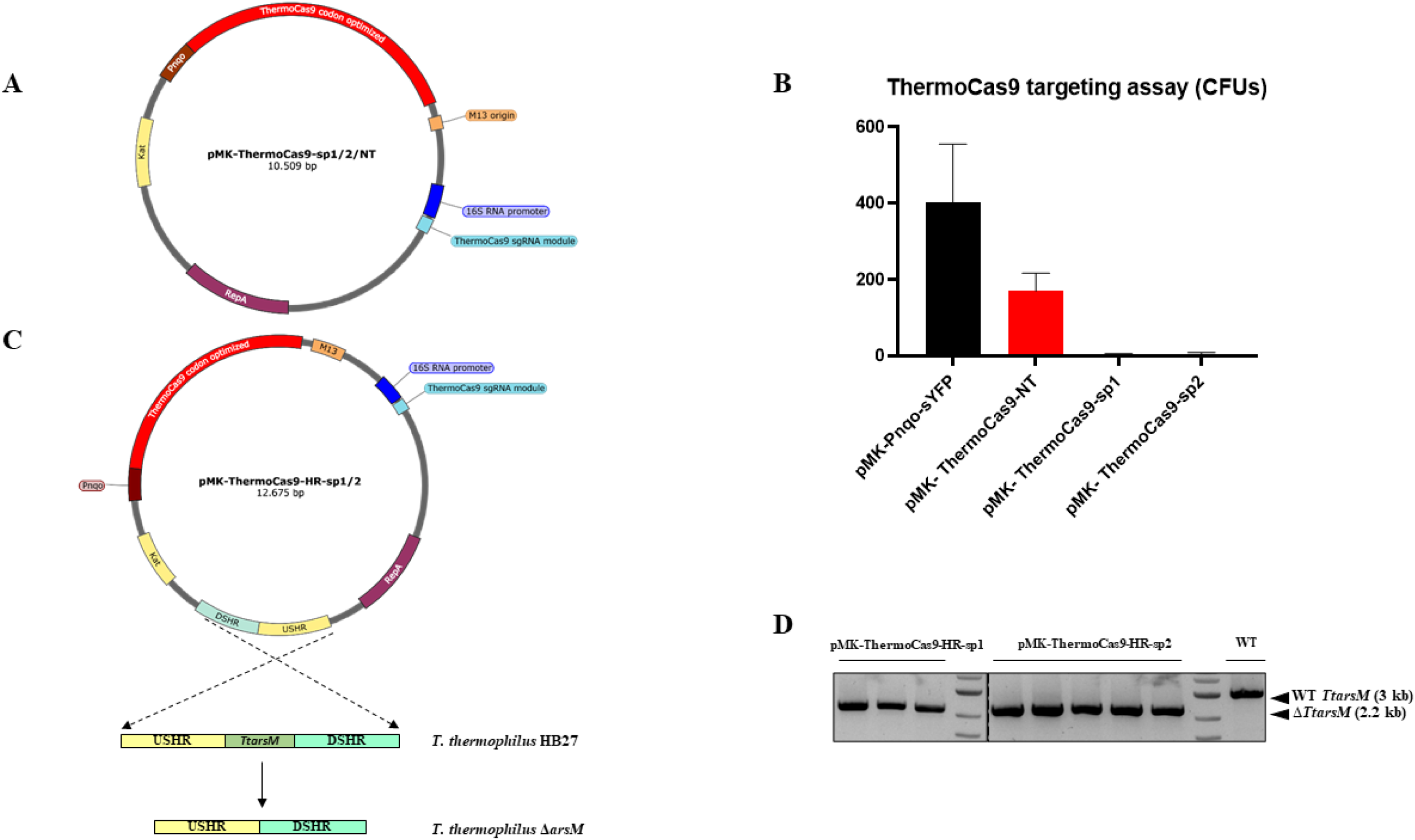
**ThermoCas9-based genome engineering in *T. thermophilus* HB27. A)** pMK-ThermoCas9-sp1/2/NT targeting vectors. **B)** Graphical representation of the ThermoCas9 targeting assay results (CFUs), for assessing the ThermoCas9 toxicity and targeting efficiency in *T. thermophilus* HB27. Average values from three biological replicates are shown, with error bars representing standard deviations. **C)** pMK-ThermoCas9-HR-sp1/2 editing vectors, employed for the genomic deletion of the *TtarsM* gene. **D)** Agarose gel electrophoresis showing the resulting products from genome-specific colony PCRs on *T. thermophilus* colonies formed from the ThermoCas9-based *TtarsM* deletion process. A wild type colony was subjected to the same PCR and the related product is shown here as negative control for *TtarsM* deletion. The expected sizes of the PCR amplification products that correspond to the wild type and Δ*TtarsM* genotypes are indicated with black arrows.

We set out to develop and test the efficiency of an HR ThermoCas9 based CS genome editing tool in *T. thermophilus* HB27. For this purpose, we introduced an HR template for the deletion of the *TtarsM* gene into the 3 previously described ThermoCas9 vectors. The HR template was composed of the fused 1kb upstream and downstream flanking regions of the *TtarsM* gene (Fig. 6 C). The three resulting editing vectors, namely pMK-ThermoCas9-HR-NT, pMK- ThermoCas9-HR-sp1 and pMK-ThermoCas9-HR-sp2, were transformed into *T. thermophilus* HB27 cells, recovered at 70°C, and grown on selective agar plates overnight at 60°C. Colony PCR with genome-specific primers was subsequently employed to screen several colonies for each transformation (Fig. S4, Table S2). None of the colonies from the pMK-ThermoCas9-NT transformation were clean Δ*TtarsM* mutants (0/10 colonies), and only a small number of colonies were mixed wild type/Δ*TtarsM* mutants (2/10 colonies) (Fig. S4 A, Table S2). On the other hand, almost all the screened colonies from the pMK-ThermoCas9-sp1 transformation were clean Δ*TtarsM* mutants (19/19 colonies) (Fig. S4 B, Table S2); most of the screened colonies from the pMK-ThermoCas9-sp2 transformation were clean Δ*TtarsM* mutants (13/18 colonies), and the remaining were mixed wild type/Δ*TtarsM* mutants (5/18 colonies) (Fig. S4 C, Table S2); the latter result suggests that less efficient ThermoCas9 targeting is obtained when employing spacer 2. Subsequently, DNA sequencing on randomly selected clean Δ*TtarsM* mutant colonies was performed to verify the correctness of the genome editing (Fig. S4 D, Table S2).

Aiming to test the temperature limit of the developed ThermoCas9-based genome-editing tool, we repeated the editing experiment increasing the plating temperature to 65°C, corresponding to the temperature limit of the pMK18 backbone for propagation. Under these conditions, the number of colonies formed upon transformation with the pMK-ThermoCas9-sp1 and pMK- ThermoCas9-sp2 was of 3 and 5 vectors, respectively, much lower compared to the corresponding numbers when the plating temperature was 60°C (Table S2). This can be ascribed to the high ThermoCas9 targeting activity at 65°C (43) and decreased vector stability at 65°C. Nonetheless, the DNA sequence of all the screened colonies confirmed that they were clean Δ*TtarsM* mutants, demonstrating the high efficiency of the developed tool at 65°C (Fig. 6 D).

Finally, assuming that the curing of the editing plasmid from a Δ*TtarsM* mutant strain would facilitate additional editing steps, we randomly selected a Δ*TtarsM* mutant colony for inoculation in liquid, antibiotic-free TM medium, for two culturing rounds at 65°C and then plated the cultures on TM agar plates with and without antibiotic. Multiple colonies were found on the antibiotic-free plate and no colonies on the plate supplemented with the antibiotic, demonstrating that the cells were cured from the edited vector. Seven of these colonies were randomly selected and the absence of the plasmid confirmed by colony PCR using *thermoCas9-*specific primers (Fig. S4 E).

Therefore, a marker-less HR-ThermoCas9-based CS genome editing tool was developed for *T. thermophilus* HB27, highly efficient at temperatures up to 65°C. Using this tool, a *T. thermophilus* Δ*TtarsM* strain was constructed in less than 10 days (including the plasmid curing process), expanding the repertoire of available genetic tools for this microorganism and considerably accelerating the required time for editing its genome. To the best of our knowledge, this is the highest temperature reported for a CRISPR-Cas9 based genome-editing tool to date.

### *Tt*ArsM mutant is more sensitive to arsenite

To compare the arsenic resistance of Δ*TtarsM* to that of wild type *T. thermophilus* HB27, both strains were grown in TM liquid medium with different arsenite and arsenate concentrations for 24-hours (Fig. 7). As expected, the arsenite resistance of Δ*TtarsM* was significantly lower than that of the wild type strain, with the corresponding MIC values being 18 mM and 40 mM, respectively (Fig. 7 A). Moreover, the resistance of the Δ*TtarsM* strain to arsenate was comparable to the wild type strain (42 mM and 44 mM, respectively) (Fig. 7 B), in agreement with its role in arsenite resistance. This result confirmed that the thermoactive arsenite methyltransferase *Tt*ArsM, is involved in arsenite detoxification and is a novel component of the arsenic resistance machinery.

**Fig. 7.**
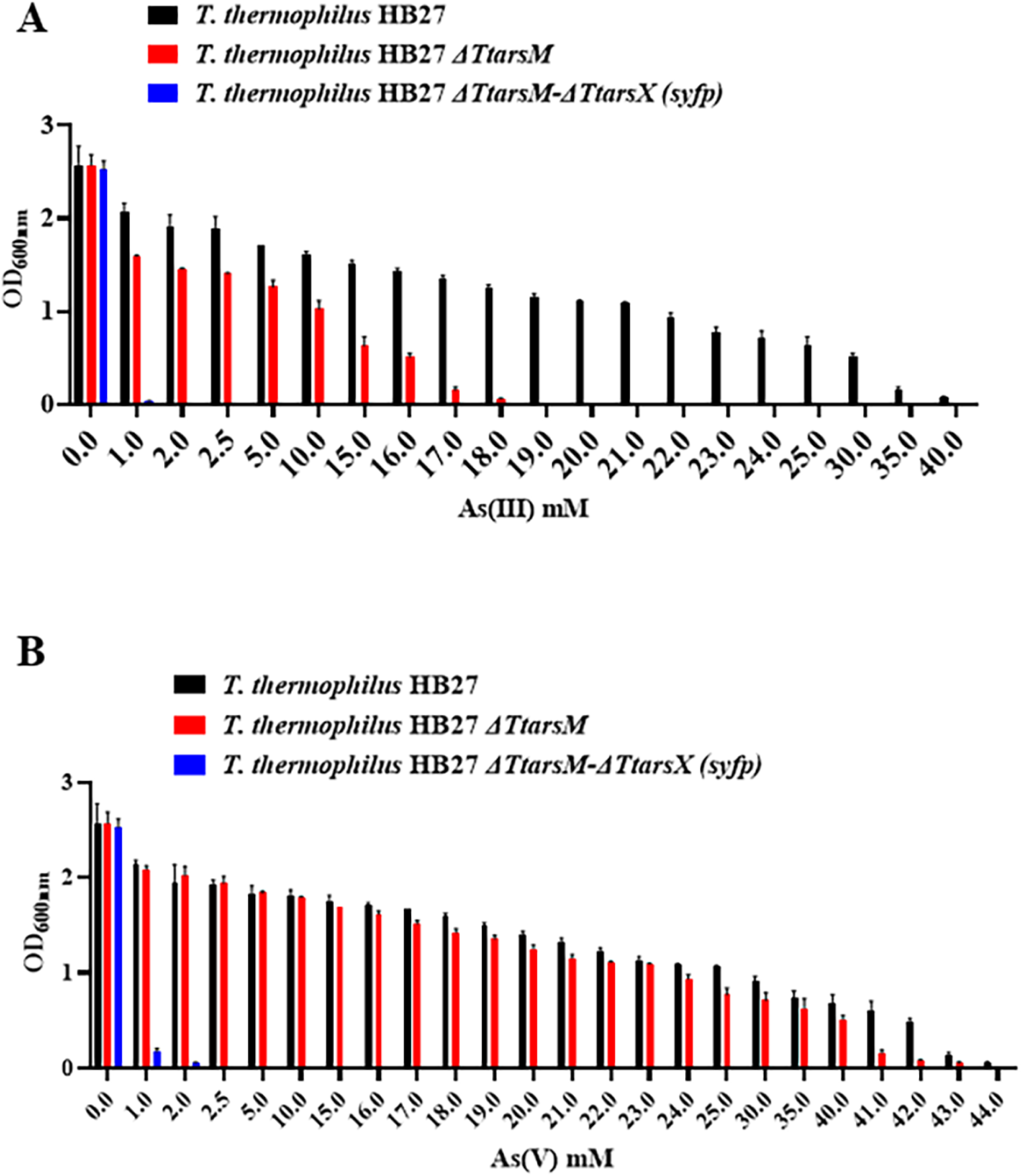
Growth of *T. thermophilus* HB27 (black), *T. thermophilus* HB27 Δ*TtarsM* (red) and *T. thermophilus* HB27 Δ*TtarsM-*Δ*TtarsX* (*syfp*) (blue) in TM medium in the presence of different concentrations of **A)** arsenite and B) arsenate measured 24 hours after inoculation. Average values from three biological replicates are shown, with error bars representing standard deviations.

### Developing a sensitive arsenic bioreporter

In a previous study, we demonstrated that *Tt*ArsX is the arsenic efflux membrane protein of *T. thermophilus* HB27 and reported that a Δ*TtarsX* mutant strain is more sensitive to arsenate and arsenite (23). In this study, we wanted to ascertain whether a strain lacking both *TtarsM* and *TtarsX* would be even more sensitive to arsenic ions than the single mutants Δ*TtarsX* and Δ*TtarsM* strains, and therefore could represent an even better bioreporter strain for arsenic detection. For this purpose, the HR ThermoCas9 based CS editing tool was employed to exchange *TtarsX* gene in the genome of the Δ*TtarsM* strain with the *syfp* reporter gene (27), setting the expression of the encoded thermotolerant yellow fluorescence protein (sYFP) under the control of the arsenic responsive *TtarsX* promoter (*p_arsX_*). The employed editing vector, denoted as pMK-ThermoCas9- HR-*syfp*, contained a spacer that targets the *TtarsX* gene (5’- TTTCGACGGAGGAGGCCTTGGCC-3’) and an HR-template composed of the 1kb upstream flanking genomic region of *TtarsX* followed by *syfp* and the 1kb downstream flanking genomic region of *TtarsX*. Ten colonies grown after transformation of pMK-ThermoCas9-HR-*syfp* vector into *T. thermophilus* Δ*TtarsM* cells were screened by colony PCR with genome-specific primers and sequenced; eight of them were clean *T. thermophilus* HB27 Δ*TtarsM-*Δ*TtarsX* (*syfp*) knock-in mutants (Fig. S5), also proving that the developed tool was highly efficient for gene insertions and substitutions.

The double mutant strain was challenged with different arsenite and arsenate concentrations in TM liquid medium. As shown in Figure 7A and 7B, arsenite resistance is strikingly lower (0.5 mM) compared to that of the wild type (40 mM), Δ*TtarsM* (18 mM) and Δ*TtarsX* strains (3 mM) (23). Interestingly, the Δ*TtarsM-*Δ*TtarsX* (*syfp*) strain showed also lower resistance to arsenate compared to the single mutant *ΔTtarsX* (2 mM and 3 mM, respectively) (22).

To evaluate the sensitivity to arsenate and arsenite of the *whole cell* bioreporter system, exponentially growing cultures of *T. thermophilus* HB27 Δ*TtarsM-* Δ*TtarsX* (*syfp*) were treated with increasing concentrations of arsenite and arsenate and the intensity of the emitted fluorescence was compared (Fig. 8). The background fluorescence of the Δ*TtarsM-*Δ*TtarsX* (*syfp*) strain was low, indicating that the system is repressed in the absence of metal ions. Moreover, the developed bioreporter system was able to detect arsenite and arsenate concentrations as low as 0.5 µM (Fig. 8). This performance substantially overtakes the detection limit of the previously developed arsenite and arsenate bioreporter system, which was based on the *T. thermophilus* Δ*TtarsX* strain and plasmid-based expression of the β-galactosidase (23).

**Fig. 8.**
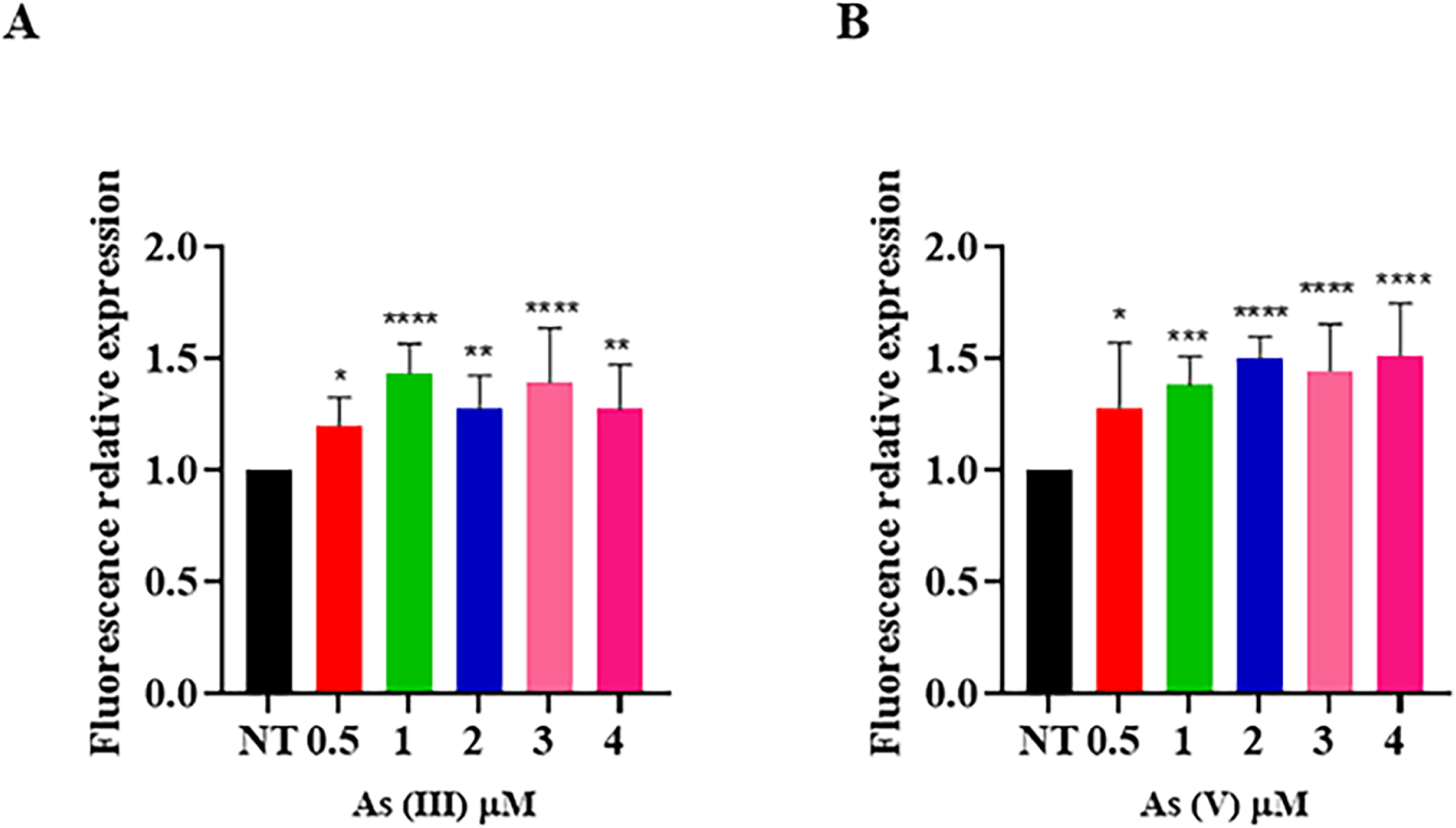
**Bioreporter activity.** *T. thermophilus* HB27 Δ*TtarsM-*Δ*TtarsX* (*syfp*) bioreporter strain challenged with increasing concentrations of **A)** arsenite and **B)** arsenate. Average values from three biological replicates are shown, with error bars representing standard deviations. Statistical analysis was performed using one-way ANOVA; significant differences are indicated as: * p < 0.05, ** p < 0.01, *** p < 0.001, **** p < 0.0001.

## Discussion

In this study, we aimed to identify novel proteins involved in the arsenic resistance system of *T. thermophilus* HB27 and employed *Tt*SmtB as a starting point looking beyond its transcriptional regulation activity. As *Tt*SmtB contains a protein interaction domain (32), we set out to identify putative *Tt*SmtB interacting proteins with a role in arsenic metabolism/detoxification, following an immunoprecipitation and comparative proteomics approach. This strategy led to the discovery of *Tt*ArsM, the first prokaryotic thermoactive arsenite SAM-dependent methyltransferase, evolutionarily distant from other known arsenite methyltransferases.

The original structure and activity mechanism of *Tt*ArsM was explored *in silico*. Like other arsenite methyltransferases known to date, *Tt*ArsM contains a C-terminal SAM-dependent methyltransferase domain and an N-terminal domain comprising only one (instead of the usually three) conserved cysteine; the other two catalytic cysteines required for arsenite coordination might be replaced by two histidines identified by docking analysis. Interestingly, H40 is part of an arsenite binding domain typical of many ArsR metalloregulatory proteins (45) and H179 is located close to the cysteine (C177) of the arsenite methyltransferase of the extremophilic alga *Cyanidioschyzon merolae* (46). Moreover, the conservation of these three amino acids among the putative arsenite methyltransferases in the *Thermus* genus supports their possible role in catalysis and suggests an adaptation in this group of microorganisms.

Indeed, the role of the cysteine and histidine residues in the structure-function of *Tt*ArsM predicted *in silico* was demonstrated by site-directed mutagenesis as the heterologous expression in *E. coli* of *Tt*ArsM mutants conferred lower arsenite resistance than *Tt*ArsM.

To date, only for a few arsenite methyltransferases, the reaction mechanism has been reported (13, 20, 36, 37, 47). Those described possess three or at least two cysteine residues present in their catalytic site that can methylate the arsenic sequentially in its trivalent form through alternating reduction and oxidative methylation reactions; noteworthy, different enzymes produce mono- di- and tri- methylated forms of arsenic in diverse amounts, highlighting a biochemical diversity in the arsenite methylation mechanism (36). The newly discovered *Tt*ArsM, highly conserved within *Thermus* genus, possesses only one cysteine, and can methylate As(III) mainly into DMAs and a smaller amount of MMAs as determined by GC-MS analysis of the products of the *in vitro* assay. To the best of our knowledge, this is the first arsenite methyltransferase functioning with a single cysteine in the active site.

The discovered transcriptional and post-translational interaction of *Tt*ArsM with the transcriptional regulator *Tt*SmtB was investigated in more detail. It was demonstrated that *Tt*SmtB binds to the promoter region of *TtarsM* and that this binding is stabilised by the *Tt*SmtB:*Tt*ArsM complex. Moreover, Co-Ip experiments confirmed the interaction of *Tt*SmtB with *Tt*ArsM and showed a reverse correlation between the stability of the complex and arsenic concentration. Presumably, the complex enhances the repression of *TtarsM* transcription in the absence of arsenic ions through a novel mechanism. Hence, through this analysis, we shed light on a novel kind of interaction, rarely described for bacteria, in which the transcriptional repressor of a gene interacts with the protein product of the gene that it regulates (41, 48).

An example of an enzyme that can modulate the transcriptional activity of regulators by protein-protein interaction is reported in the cysteine metabolism of *Bacillus subtilis* where the stable complex formed by CymR (the master regulator of the system) and CysK (*O*-acetyl-L-serine-thiol-lyase) represses the transcription of the genes involved in the cysteine pathway (including *cysK* gene itself) when cysteine concentration is low. The advantage of this regulatory mechanism is that it employs enzymes that can specifically recognize their substrates or allosteric effectors; thus, enzymes and/or transcriptional regulators can act simultaneously as intracellular molecular sensors and participate to their own transcriptional regulation (40).

The role of *Tt*ArsM in *T. thermophilus* HB27 arsenic detoxification was also demonstrated *via* the construction and characterization of a *ΔTtarsM* mutant strain. For this purpose, a marker-free, homologous recombination and ThermoCas9 based counter-selection genome editing tool was developed, which was highly efficient and active at temperatures up to 65°C. Our tool equals the highest reported temperature for a Cas9-based editing tool to date (49, 50). The characterization of the *T. thermophilus ΔTtarsM* strain confirmed its expected higher sensitivity to arsenite, but not arsenate, compared to the wild type strain.

To better define the role of *Tt*ArsM in the context of the already characterized components of the arsenic resistance system, a double mutant was constructed upon exchanging the *TtarsX* efflux pump gene in the *T. thermophilus* Δ*TtarsM* genome with the gene encoding the yellow fluorescent protein; the double mutant resulted much more sensitive to arsenite and arsenate treatment. Hence, it was demonstrated that *Tt*ArsM and *Tt*ArsX are critical players of the arsenite detoxification system. This is the first example of a successfully insertion of a heterologous gene on the *T. thermophilus* genome by genome editing. The double mutant strain was also considered as a sensitive bioreporter for the development of a *whole cell* biosensor system. Indeed, it was able to detect arsenite and arsenate concentrations as low as 0.5 µM, showing 40 times higher sensitivity when compared to the previously developed *T. thermophilus* HB27 *ΔTtarsX*- plasmid-based biosensor (23).

In conclusion, this study explores a unique strategy to identify novel enzymes and/or regulative networks in non-model bacteria and expands the repertoire of genetic systems for hyperthermophiles. In addition, this work has resulted in a gain of insight into the arsenite/arsenate detoxification mechanism, particularly in that of *T. thermophilus*. On top of that, this has allowed us to develop a highly robust and sensitive biosensor.

## Materials and Methods

### *T. thermophilus* HB27 cell-free extract preparation

*T. thermophilus* HB27 cultures were grown aerobically at 70°C in TM medium, as previously described (24). Once the cultures reached 0.5 OD_600nm_, they were treated either with 8 mM NaAsO_2_ or with 12 mM NaH_2_AsO_4_ (Sigma) [the used concentrations were below the previously reported MIC values for arsenate and arsenite (22, 23)] or they remained untreated. Samples were harvested from each culture, either immediately after treatment or 60 min post-treatment. The samples were centrifuged, the precipitates were resuspended in phosphate buffer (20 mM Na_3_PO_4_ pH 7.5) supplemented with protease inhibitor cocktail (Thermo Scientific) and the resuspended cells were lysed by sonication (10 cycles: 30’’on/30’’off, power 40%, Misonix® Sonicator Ultrasonic Processor XL). The lysates were centrifuged and the cell-free extracts (CFE) used for pull-down assays.

### Purification of recombinant *Tt*SmtB, Immobilized Metal Affinity Chromatography (IMAC) and pull-down

C-terminal His-tagged *Tt*SmtB was purified from *E. coli* BL21-CodonPlus (DE3)-RIL cells transformed with the pET28/*Ttsmt*B vector, as previously described (22). Purified C-terminal His-tagged *Tt*SmtB (2 mg), was incubated with 200 µL of Ni^2+^-NTA resin (Sigma-Aldrich) equilibrated in 20 mM Na_3_PO_4_, 0.5 M NaCl, 20 mM imidazole, pH 7.5 for 16 h at 4°C and then washed three times with the same buffer to remove unbound proteins. *T. thermophilus* HB27 CFE, treated with arsenite, or treated with arsenate, or not treated were incubated with the functionalized resin (Ni^2+^-NTA/*Tt*SmtB) for 16 h at 4°C under stirring conditions. Subsequently, the resin was extensively washed and the interacting proteins were eluted with 20 mM Na_3_PO_4_, 0.5 M NaCl, 0.5 M imidazole; pH 7.5. As negative controls, samples of Ni^2+^-NTA resin not functionalized with *Tt*SmtB were incubated with the same *T. thermophilus* HB27 CFE.

### *In situ* hydrolysis and LC-MS/MS analysis

The fractions eluted from the pull-down process were analyzed by 15 % SDS-PAGE and *in situ* hydrolyzed for mass spectrometry analysis. Specifically, mono-dimensional SDS-PAGE gel was coloured with Coomassie Brilliant Blue; the revealed bands were cut and de-stained with 100 µL of 0.1 M ammonium bicarbonate (AMBIC) and 130 µL of acetonitrile (ACN). Each band was hydrolyzed *in situ* with 0.1 µg/µL trypsin in 10 mM AMBIC, incubated at first for 1.5 h at 4°C and then for an additional 16 h at 37°C. The hydrolysis reactions were stopped by adding acetonitrile and 0.1% formic acid; then, the samples were filtered and dried in a Savant vacuum centrifuge before being analyzed by LC-MS/MS mass spectrometry. In detail, before analysis, the samples were dissolved in 10 µL of 0.1% formic acid, and 5 µL were directly loaded into the instrument. Reverse-phase capillary liquid chromatography (HPLC 1200 system experiments), followed by MS analysis, was performed using a binary pump system connected to a nano-spray source of the mass spectrometer (28, 51). The latter is represented by a hybrid Q-TOF spectrometer (MS CHIP 6520 QTOF) equipped with a chip (Agilent Technologies).

### *In silico* analysis

Analysis of the LC-MS/MS data, using Mascot software (http://www.matrixscience.com/search_form_select.html) allowed the identification of putative *Tt*SmtB-interacting proteins. Among these proteins, TTC0109 (UniProt code Q72LF0), herein named *Tt*ArsM, was further analyzed using the UniProt database (http://www.uniprot.org); homologous proteins and conserved domains were identified by performing a Blast analysis (https://blast.ncbi.nlm.nih.gov/Blast.cgi).

Phylogenetic tree of archaeal and bacterial arsenite methyltransferases, SAM-dependent methyltransferases and methyltransferase domain-containing proteins, including *Tt*ArsM from *T. thermophilus* HB27, was conducted in MEGA X (52). The amino acidic sequences used for the construction of the phylogenetic tree are *Tt*ArsM from *T. thermophilus* HB27, arsenite methyltransferase from *Rhodopseudomonas palustris* (13), *Methanosarcina acetivorans*, *Clostridium* sp BMX (53), *Halobacterium salinarum* (54), *Pseudomonas alcaligenes* (36)*, Cyanidioschyzon merolae* (46); a SAM-dependent methyltransferase from 5 members of the *Thermus* genus (*islandicus, caldilimi, antranikianii, oshimai* and *brockianus*); a SAM-dependent methyltransferase from *Mesorhizobium amorphae*, *Anaerolineae bacterium*; a methyltransferase domain-containing protein from *Sanguinobacter* sp. Phylogenetic reconstruction was accomplished using the maximum likelihood statistical method.

The alignment of *Tt*ArsM to its templates was based on a multiple sequence alignment, performed with the program Clustal Omega (55); the amino acidic sequences used for the construction of the alignment of functionally characterized archaeal and bacterial arsenite methyltransferases are: *Tt*ArsM from *T. thermophilus* HB27, 5 members of the *Thermus* genus (*islandicus, caldilimi, antranikianii, oshimai* and *brockianus*), *Clostridium* sp BMX, *R. palustris*, *M. acetivorans*, *H. salinarum*, *P. alcaligenes* and *Cyanidioschyzon merolae* (46).

Models of *Tt*ArsM were generated through I-TASSER (56) (https://zhanglab.ccmb.med.umich.edu/I-TASSER/) using as input the complete sequence of *Tt*ArsM (C-score = -2.5). The dimeric structure was predicted using the GalaxyWEB tool (http://galaxy.seoklab.org/index.html) (57). The molecular dockings of *Tt*ArsM with arsenite and SAM were generated using the Hex Protein Docking server (58). 100 rigid-body docking solutions were generated per case and the best 10 were refined by energy minimization. The proposed model for the metal ion docked into *Tt*ArsM is the structure with the smallest distance between arsenite-histidine and cysteine-SAM (4.33 Å from H40 and 5.77 Å from H179 in *Tt*ArsM model and 4.40 Å from C77).

### Cloning, expression and purification of recombinant *Tt*ArsM and *Tt*ArsM mutants

The pET30b(+)*/TtarsM* vector was constructed for the expression and subsequent purification of the C-terminal His-tagged version of *Tt*ArsM. For the construction of the pET30b(+)*/TtarsM*, *TtarsM* gene was PCR amplified from *T. thermophilus* HB27 genome, using Taq DNA polymerase (Thermo Fisher Scientific) and primers containing the NdeI (*arsMfw,* Table S3) and HindIII (*arsMrv,* Table S3) restriction sites at their 5’-ends. The PCR product was purified, digested with the NdeI and HindIII restriction enzymes (NEB), and ligated (T4 ligase, NEB) into NdeI/HindIII-digested pET30b(+) vector (Novagen). The ligase mixture was transformed into *E. coli* TOP10F’ cells were plated on LB agar plates supplemented with 50 µg/mL kanamycin (Sigma-Aldrich). Single colonies were selected and inoculated in LB liquid medium supplemented with 50 µg/mL kanamycin. Plasmid isolation and sequencing were subsequently performed before transforming *E. coli* BL21-CodonPlus (DE3)- RIL cells with pET30b(+)*/TtarsM* vector.

To obtain mutation of *TtarsM* gene sequence at specific sites, the QuickChange II-E Site-Directed Mutagenesis Kit (Agilent Technologies) was employed; pET30b(+)*/TtarsM* was used as a template and amplified with three different mutagenic primer pairs (Table S3) to get pET30b(+)*/TtarsM* C77S, pET30b(+)*/TtarsM* H40A and pET30b(+)*/TtarsM* H179A vectors. The reaction mixtures were transformed into *E. coli* TOP10F’ cells were plated on LB agar plates supplemented with kanamycin (50 µg/mL). Single colonies were randomly selected and inoculated in LB liquid medium supplemented with kanamycin (50 µg/mL). Plasmid isolation was subsequently performed and *E. coli* BL21- CodonPlus (DE3)-RIL cells were transformed with sequence-verified pET30b(+)*/TtarsM* C77S, pET30b(+)*/TtarsM* H40A and pET30b(+)*/TtarsM* H179A vectors.

For protein expression of the His-tagged versions of *Tt*ArsM C77S, *Tt*ArsM H40A and *Tt*ArsM H179A catalytic mutants, the recombinant *E. coli* BL21- CodonPlus (DE3)-RIL strains were cultured in LB medium supplemented with kanamycin (50 µg/mL) and chloramphenicol (33 µg/mL). Protein expression was induced via the addition of 1 mM isopropyl-1-thio-β-D-galactopyranoside (IPTG) when the cultures reached 0.7 OD_600nm_. The cultures were further incubated with vigorous shaking at 37°C for 16 h, then centrifuged, resuspended in lysis buffer (20 mM NaP pH 7.4, 50 mM NaCl and 20 mM imidazole) supplemented with protease inhibitor cocktail (Thermo Scientific) and lysed by sonication (10 cycles: 30’’on/30’’off, power 40%, Misonix® Sonicator Ultrasonic Processor XL). The lysates were centrifuged and the supernatants used for the purification on HisTrap HP columns (1 mL; GE Healthcare) connected to an AKTA Explorer system (GE Healthcare). The fractions containing His-tagged *Tt*ArsM proteins were eluted from the columns using a linear gradient of the elution buffer (20 mM NaP pH 7.4, 50 mM NaCl and 500 mM Imidazole). The eluted protein fractions were subjected to SDS-PAGE analysis and the fractions containing purified *Tt*ArsM were pooled and dialyzed for 16 h at 4°C in 20 mM NaP pH 7.4 buffer supplemented with protease inhibitor cocktail (Thermo Scientific). The identity of the purified His-tagged *Tt*ArsM protein was confirmed by mass spectrometry and protein aliquots were stored at -20°C.

### *Tt*ArsM quaternary structure assessment

The native molecular mass of *Tt*ArsM was determined by loading 500 µg of the purified protein onto an analytical Superdex PC75 column (3.2 by 30 cm) connected to an AKTA Pure system, in 50 mM Tris-HCl, pH 7.5, 0.2 M KCl buffer. The column was calibrated using a set of gel filtration markers (low range, GE Healthcare), including Ovalbumin (43.0 kDa), Carbonic anhydrase (29.0 kDa), RNase A (13.7 kDa) and Aprotinin (6.5 kDa) as previously described (24).

### Methyltransferase activity assay

According to the manufacturer’s protocol, the *Tt*ArsM arsenite methyltransferase activity was measured using the SAM510: SAM Methyltransferase Assay Kit (G-Biosciences) with modifications regarding the temperature, the SAM concentration and the reaction time (33–35). The assay relies on the degradation of S-adenosylhomocysteine (SAH) into urate and hydrogen peroxide by a mixture of enzymes (adenosylhomocysteine nucleosidase, adenine deaminase, xanthine oxidase). Then, the reaction of hydrogen peroxide with 4- aminoantipyrine produces 5-dichloro-2-hydroxybenzene sulfonic acid (DHBS) with ε_mM_=15.0 at 510_nm_. A typical reaction mixture containing: 200 µM As(III), 800 µM SAM, 3.1 μM of the enzyme, SAM enzyme mixture and SAM colorimetric mix in a final reaction volume of 115 µL, was incubated for 1h at 50°C in a Synergy™ HTX Multi-Mode Microplate Reader (BioTek). The same reaction mixture was tested with 10 µg of *Tt*ArsC or 10 µg of *Tt*SmtB, as negative controls. One unit of arsenite methyltransferase produces 1.0 µmol of DHBS per minute at 50°C under the conditions described above. Preliminary assays were performed to define substrate saturating concentrations, varying the As(III) and SAM concentrations from 50 µM to 300 µM and from 200 µM to 1.2 mM, respectively.

### In vitro arsenite methylation

As(III) methylation by *Tt*ArsM was determined in an assay solution containing 10 µM *Tt*ArsM, 250 µM As(III), 6 mM glutathione (GSH) and 1 mM SAM, in Na-phsphate 50 mM, pH 7.4 at 65°C for 24 h; the same reaction without *Tt*ArsM was used as the negative control (control sample). The reactions were terminated by the addition of 10% (v/v) H_2_O_2_. The oxidized solution samples were filtered through 0.22 µm MCE syringe filters and used for GC-MS analysis.

For the detection of the methylated products, GC analyses were performed using Agilent GC 6890, coupled with a 5973 MS detector.

200 µl of the control sample and the enzyme mixture were treated with 800 µL of methanol (Sigma-Aldrich). The supernatants were recovered and dried under vacuum. The samples were solubilized in methanol (10 µl) and 1 µl was analyzed by GC-MS. The column used was an HP5 capillary (30 m × 0.25 mm, 0.25 mM, 5% polisilarilene 95% PDMS). Helium was used as the carrier gas at a rate of 1.0 mL min^−1^. The GC injector was maintained at 230 °C, while the oven temperature was held at 40 °C for 5 min and then increased to 280 °C at 20 °C/min and held for 5 min for a total separation time of 20 min. The analyzer temperature was kept at 250 °C. The collision energy was set to a value of 70 eV, and fragment ions generated were analyzed mass range 20–450 m/z. The identification of each compound was based on the combination of retention time and fragmentation spectra matching those collected into the NIST 05 Mass Spectral Library. The identification was reliable when the matching values were higher than 700 according to the NIST guidelines (59). The analyses were performed in triplicate for each sample.

### Co-Ip assay

Protein-protein interaction between *Tt*SmtB and *Tt*ArsM was *in vitro* verified via co-immunoprecipitation assays that employed recombinant *Tt*SmtB, recombinant His-tagged-*Tt*ArsM, anti-*Tt*SmtB antibodies (GeneCust™) and His-tag antibodies (Sigma-Aldrich). His-tag removal from recombinant *Tt*SmtB was performed as previously described (22).

A typical Co-Ip mixture contained 1 mL of Co-Ip buffer (50 mM Tris-HCl, pH 7.5, 150 mM NaCl, 10% glycerol and 0.1% Triton X-100), 5 µg of *Tt*SmtB, 5 µg of His-tagged *Tt*ArsM and was incubated at 4°C for 2h in continuous rotation. In some cases, arsenite, arsenate, cadmium and antimony at 1:0, 1:25, 1:50, 1:100 molar ratios pre-incubated with *Tt*SmtB for 10 min at 60°C were added. As controls, *Tt*ArsM (5 µg), *Tt*SmtB (5 µg) and *Tt*GalA (38) were also separately incubated under the same conditions. All the samples were subjected to immunoprecipitation using 2 µL of purified anti-*Tt*SmtB antibodies (2 µg/µL) (GeneCust™) for 3 h at 4°C in continuous rotation, before adding 15 µL of Protein A-Sepharose beads (Sigma-Aldrich) and allowing the incubation to continue for further16 h at 4°C. The formed immunocomplexes were washed with Co-Ip buffer and analyzed by Western Blot on 15% SDS-PAGE, using PVDF membranes (Millipore) and anti-poly-Histidine-Peroxidase antibodies (Sigma-Aldrich) diluted 1:10000, as previously described (60).

The ImageJ software (https://imagej.net/) was used for densitometric analysis of the formed bands, setting as maximum value the intensity of the band of the sample containing *Tt*ArsM and *Tt*SmtB in the absence of any metal. Each experiment was performed in three technical and two biological replicates; statistical analysis was performed using one-way ANOVA and significant differences are indicated as * *p* < 0.05, ** *p* < 0.01, *** *p* < 0.001.

### Electrophoretic mobility shift assay (EMSA)

Electrophoretic mobility shift assays (EMSA) were performed to determine the *in vitro* binding of *Tt*SmtB to the promoter region upstream *TtarsM*. The 108 bp chromosomal region that encompasses the start codon of the *TtarsM* gene and the 105 bp-long region upstream, denoted as *p_arsM_*, was PCR amplified with Taq DNA polymerase (Thermo Fisher Scientific) using the *p_arsM_ Fw* and *p_arsM_ Rv* primers (Table S3). EMSA reactions were set up as previously described (23, 61) in the presence of 1µg of poly (dI-dC), 20 ng *p_arsM_* and increasing concentrations of *Tt*SmtB (1, 2, 3, 5 and 10 µM, considering *Tt*SmtB as a dimer) using SYBR™ Gold Nucleic Acid Gel Stain for band detection. The EMSA reactions that simultaneously employed *Tt*SmtB and *Tt*ArsM were set up in identical conditions using 3 µM of each protein alone or in combination.

### Arsenic tolerance of *E. coli* expressing *Tt*ArsM

The following strains were inoculated in 10 mL of LB pre-cultures, supplemented with kanamycin (50 µg/mL) and chloramphenicol (33 µg/mL): i) *E. coli* BL21- CodonPlus (DE3)-RIL: pET30*/TtarsM,* ii) *E. coli* BL21-CodonPlus (DE3)-RIL: pET30*/ TtarsM C77S,* iii) *E. coli* BL21-CodonPlus (DE3)-RIL: pET30*/ TtarsM H40A,* iv) *E. coli* BL21-CodonPlus (DE3)-RIL: pET30*/ TtarsM H179A* and v) *E. coli* BL21-CodonPlus (DE3)-RIL: pET30 (control). The pre-cultures were incubated at 37°C for 16 h at 180 rpm. Subsequently, 50 mL LB cultures, supplemented with antibiotics, were inoculated with the pre-cultures to initial 0.08 OD_600nm_ and incubated at 37°C and 180 rpm until 0.6 OD_600nm_ (exponential growth). At that point, protein expression was induced with 1mM IPTG and the cultures were incubated at 37°C and 180 rpm for 3 additional hours. From these growing cells, fresh LB cultures were inoculated to 0.05 OD_600nm_ and distributed to 24-well plates (1 mL per well) containing LB medium supplemented with 1 mM IPTG, kanamycin (50 µg/mL), chloramphenicol (33 µg/mL) and increasing concentrations of arsenate and arsenite (from 2.5 mM to 7.0 mM). The minimal inhibitory concentrations (MIC) endpoint for each strain were determined as the lowest concentration of arsenite at which there was the difference between grown and start culture lower than 0.01 OD_600nm_ after 16 hours of incubation at 37°C (22). All the cells up to the MIC value were able to grow if reinoculated in an arsenic-free medium. The reported values are the average of three biological replicates.

### ThermoCas9 editing and targeting constructs

The plasmids used for the ThermoCas9-based targeting and editing experiments are listed in Table S4. The vector pMK18 was used as the template for the construction of the ThermoCas9-based targeting and editing plasmids and the employed primers, the DNA templates, and the DNA fragments, which are listed in Table S4. The *thermoCas9* gene was codon harmonized, according to *T. thermophilus* HB27 codon-usage using the Galaxy/Codon harmonizer online tool (62), and it was synthesized (Twist Bioscience) (Table S5). The DNA fragments were designed with appropriate overhangs for NEBuilder HiFi DNA assembly (NEB) and they were obtained through PCR with Q5 Polymerase (NEB). The PCR products were subjected to 1% agarose gel electrophoresis and they were purified using a Zymogen gel DNA recovery kit (Zymo Research). The assembly reactions were transformed to chemically competent *E. coli* DH5α cells (NEB) and the cells were plated on LB agar plates supplemented with kanamycin (50 µg/mL). Single colonies were inoculated in LB medium supplemented with kanamycin (50 µg/mL) for overnight incubation at 37°C. Plasmid material was isolated using the GeneJet plasmid miniprep kit (Thermo Fisher Scientific), sequence verified (GATC-biotech) and 300 ng of each plasmid (pMK- ThermoCas9-NT/sp1/sp2 and pMK-ThermoCas9-HR-NT/sp1/sp2) was transformed to either *T. thermophilus* HB27 cells (22), as indicated per experimental process.

The obtained plasmid, pMK-ThermoCas9-NT, was used as the backbone to construct a new plasmid to obtain the deletion of the *TtarsX* gene (Table S4) and the insertion of the gene coding sYFP (27). The obtained plasmid (pMK- ThermoCas9-HR-syfp) was used to transform *T. thermophilus* HB27 *ΔTtarsM* cells as already described to obtain the strain denoted Δ*TtarsM*-Δ*TtarsX* (*syfp*).

### Arsenic tolerance of *T. thermophilus* HB27 wild type and mutant strains

Exponentially growing pre-cultures of *T. thermophilus* HB27 (control), *T. thermophilus* HB27 Δ*TtarsM* and *T. thermophilus* HB27 Δ*TtarsM-*Δ*TtarsX (syfp)* were diluted to 0.08 OD_600nm_ in 10 mL TM cultures containing increasing concentrations of arsenite and arsenate (from 0.1 mM to 50 mM). The cultures were incubated aerobically at 70°C for 18 h and the MIC values were determined as the lowest concentrations of arsenite and arsenate that completely inhibited the growth of a strain (22). The reported values are the average of three biological replicates.

### Bioreporter activity measurement

Overnight cultures of *T. thermophilus* HB27 Δ*TtarsM*-Δ*TtarsX* (*syfp*) strain were diluted to 0.08 OD_600nm_ in TM medium and then grown aerobically at 70°C until 0.5 OD_600nm_. The cultures were divided into samples of 5 mL each and subsequently supplemented with increasing concentrations of arsenite and arsenate (0.5 μM to 4 μM). After 1-hour of incubation at 70°C, 200 μL of each cell sample were removed and centrifuged for 5’ at 6000 rpm. The pellets were washed twice with equal volumes of PBS 1X and resuspended with equal volumes of PBS 1X before being distributed into a 96-well plate. sYFP fluorescence intensity of each sample was measured employing a Synergy™ HTX Multi-Mode Microplate Reader (BioTek), using excitation and emission wavelengths of 458 nm and 540 nm, respectively. The measured fluorescence intensities were normalized for the optical density of each sample at 600 nm. The measured fluorescence was reported as fluorescence relative expression, assuming that the fluorescence value of not treated cells (control) was 1.

Each experiment was performed in three technical and biological replicates. Statistical analysis was performed using one-way ANOVA; significant differences are indicated as: * *p* < 0.05, ** *p* < 0.01, *** *p* < 0.001.

## Acknowledgements

The authors kindly acknowledge Prof. Josè Berenguer from Universidad Autònoma de Madrid for kindly providing the pMK-Pnqo-syfp vector. Prof. Lucia Cavalca and Prof. Gian Attilio Sacchi from the University of Milan also acknowledged for donating disodium methyl arsonate hexahydrate (MMAs) (ChemService) and dimethylarsinic acid (DMAs) (Merck Life Science S.r.l.). The authors also acknowledge Prof. Angela Amoresano (Department of Chemical Sciences, University of Naples Federico II) for helpful discussion. Acknowledgements go to grant ERA-NET Cofund MarTERA, who funded the project: “FLAshMoB: Functional Amyloid Chimera for Marine Biosensing”. This project has received funding from the European Research Council (ERC), under the European Union’s Horizon 2020 research and innovation programme, grant agreement No. 834279, and from the Netherlands Organization for Scientific Research (NWO, Spinoza grant SPI 93-537) to JvdO.

## Supplemental Tables

**TABLE S1.** List of *Tt*SmtB cytosolic interactors.

**TABLE S2.** Results from genome-specific colony PCRs on *T. thermophilus* HB27 colonies formed from the ThermoCas9-based *TtarsM* deletion experiments.

**TABLE S3.** List of the primers used in this work.

**TABLE S4.** List of the PCR products used for the HiFi assembly reactions to construct the ThermoCas9-based targeting and editing plasmids. The primers and templates used for the PCR reactions are also included in this list.

**TABLE S5.** The sequence of the synthesized *thermoCas9* gene, which is codon-harmonized according to the codon usage of *T. thermophilus* HB27.

## Supplemental Figure captions

**Fig. S1 TTC0109 3D model (wild type and mutants) and docking with arsenite and SAM** (the arsenic atom is the purple sphere, the oxygen atoms are the red spheres, SAM is orange). The H40 and H179 residues coordinating arsenite are coloured blue. The C77 residue is coloured black**; B)** TTC0109 C77S; **C)** TTC0109 H40A; **D)** TTC0109 H179A, blue the mutated position.

**Fig. S2 Purification of recombinant *Tt*ArsM. A)** SDS-PAGE analysis Lane M, protein marker. Lane NI, protein extract from non-induced cells. Lane I, protein extract from induced cells. Lane P, pure protein after His-trap chromatography. **B)** Size exclusion chromatogram of *Tt*ArsM; in the box the calibration curve. The chromatogram shows a peak corresponding to the *Tt*ArsM dimeric form. **C)** Histograms represented the average peak areas for MMAs and DMAs in *Tt*ArsM enzymatic reaction sample. These peak area values correspond to a concentration lower than the LOQ assessed to 1 mg/L by using standard molecules. The CV% values obtained were lower than 15 %.

**Fig. S3 Purification of recombinant *Tt*ArsM mutants.** SDS-analysis. Lane M, protein marker. Lane NI, protein extract from non-induced cells. Lane I, protein extract from induced cells. Lane W, unbound proteins after His-trap chromatography. Lane P, pure protein after His-trap chromatography. **A)** Recombinant *Tt*ArsM C77S. **B)** Recombinant *Tt*ArsM H40A. **C)** Recombinant *Tt*ArsM H179A.

**Fig. S4** Agarose gel electrophoresis from genome-specific colony PCRs on *T. thermophilus* HB27 colonies formed from the ThermoCas9-based *TtarsM* deletion process upon transformation with **A)** the pMK-ThermoCas9-HR-NT control-editing vector, **B)** the pMK-ThermoCas9-HR-sp1 editing vector, and **C)** the pMK-ThermoCas9-HR-sp2 editing vector. Wild type colonies were subjected to the same PCR and the related products are shown as negative controls for *TtarsM* deletion. The expected sizes of the PCR amplification products that correspond to the wild type and Δ*TtarsM* genotypes are indicated with arrows. **D)** Sequencing chromatogram of the PCR amplified *TtarsM* genomic region from a randomly selected colony, previously PCR screened as Δ*TtarsM.* **E)** Agarose gel electrophoresis after the curing process of the editing vectors shows the absence of products using pMK18-specific primers for colony PCRs on *T. thermophilus* HB27 Δ*TtarsM* colonies. A wild type colony containing the pMK18 vector was subjected to the same PCR and the related product is shown here as a negative control of the curing process.

**Fig. S5** Agarose gel electrophoresis showing the resulting products from genome-specific colony PCRs on *T. thermophilus* HB27 Δ*TtarsM-*Δ*TtarsX (syfp)* colonies formed from the ThermoCas9-based substitution process of the *TtarsX* gene by the *syfp* gene. A wild type colony was subjected to the same PCR and the related product is shown as a negative control for *TtarsX* substitution. The expected sizes of the PCR amplification products that correspond to the wild type and Δ*TtarsX* (*syfp*) genotypes are indicated with black arrows. Eight out of the ten screened clones were *T. thermophilus* Δ*TtarsM-*Δ*TtarsX* (*syfp*) knock-in mutants.

